# Lipid Nanoparticles Enable mRNA Delivery to Diverse Cell Types of the Inner Retina

**DOI:** 10.1101/2025.11.21.689792

**Authors:** Sumit Biswas, Eleonora Carpentiero, Nermina Xhaferri, Paula Streckenbach, Vijay Renigunta, Moritz Lindner

## Abstract

Lipid nanoparticles (LNPs) have emerged as a promising platform for retinal genetic therapy, offering a non-viral alternative to adeno-associated viruses (AAVs). While LNPs can transfect outer retinal cells, their tropism for inner retinal cell types remains insufficiently characterized. Here, we systematically assessed cellular tropism of conventional LNPs encapsulating chemically modified mRNA encoding mCherry in murine retinal explants and dissociated retinal cells. We compared quasi-subretinal and quasi-intravitreal administrations and evaluated how retinal degeneration and inner limiting membrane (ILM) integrity influence LNP-mediated transfections. We observed that LNPs efficiently transfected Müller glia under all experimental conditions. In addition, LNPs transfected several other retinal cell types, including neurons in dissociated cells and explants, and vascular cells exclusively in explants. Subretinal delivery resulted in significantly higher transfection rates than intravitreal administration, and overall efficiency was higher in degenerate as compared to non-degenerate healthy retinas. In healthy retinas, removal of ILM-associated barriers significantly increased transfection efficiency following intravitreal administration. Together, these findings demonstrate that conventional LNPs can transfect a broader range of retinal cell types than previously recognized and highlight LNPs as a versatile tool for mRNA delivery to the retina, with applications in gene supplementation, gene editing, and regenerative therapies for inner retinal disorders.

## Introduction

Over the last two decades the eye has emerged as a key target organ for gene therapy and numerous retinal disorders are currently the focus of preclinical and clinical development^1,2^. Voretigene neparvovec for Leber’s congenital amaurosis and revakinagene taroretcel for Macular Telangiectasia Type II have now received regulatory approval^1,3^. The immune-privileged status of the retina and the anatomical compartmentalization of the eye, the later supporting the achievement of high local vector concentrations, have both facilitated this rapid developement^4^. Although Adeno-associated viruses (AAV) have become the preferred vector for retinal gene therapy, their limited packaging capacity, makes them unideal for delivering larger genes^5^. Moreover, AAV-mediated gene therapy results in persistent transgene expression, which is undesirable for certain applications like CRISPR mediated gene editing^6^.

In recent years, lipid nanoparticle (LNP)-mediated RNA delivery has emerged as a novel variant of genetic therapy. This approach can be used when transient delivery of genetic material (mostly in form of mRNA) is desired, as exemplified by the approved mRNA-based vaccines against SARS-COV-2^7^. By delivering regulatory RNA, as in the approved Patisiran treatment for hereditary amyloidosis^8^, or mRNA encoding for gene editors, LNPs can be used acting on gene level and even as a form of classical gene therapy^9^. Moreover, the higher payload offered by LNPs makes them an ideal alternative in scenarios where the packaging capacity of AAV is insufficient^10^.

Until now, only few studies have addressed the use of LNP mediated RNA transfer in the context of the retina and retinal disease^11–16^. Early investigations demonstrated that conventional lipid nanoparticles, depending on the route of delivery, either mediate transfection primarily of the retinal pigment epithelium (RPE) cells or Müller glia following subretinal or intravitreal delivery, respectively^11,13^. More recently, optimizations in LNP formulations enabled the transfection of RPE cells also by intravitreal delivery^11^ and the design of peptide-guided LNPs as well as the introduction of certain lipid moieties have enabled efficient transfection of photoreceptors^14,16,17^. While most of these studies have been conducted in mice, there is now evidence that similar transfection patterns can be obtained in non-human primates and on cultured human donor retinas^12,16^.

Despite this progress, a comprehensive understanding of the cellular tropism of LNPs, specifically within the inner retina, is missing. In particular, it remains unclear which retinal cell types, besides Müller glia, can be transfected by conventional LNPs and how structural barriers such as the inner limiting membrane (ILM) and pathological changes associated with retinal degeneration modulate delivery efficiency. A better understanding of these aspects, however, would help to develop LNP-based treatments for a spectrum of retinal diseases including optic neuropathies, congenital stationary night blindness and age-related macular degeneration.

In this study, we systematically assess the transfection patterns of conventional LNPs in murine retinal explants. We evaluate the influence of delivery route, ILM integrity, and retinal degeneration on LNP-mediated transfection efficacy, and demonstrate that LNP tropism is broader than generally thought. Specifically, we observe that they are capable of transfecting inner retinal cell types beyond phagocytosing cells, including vasculature as well as neurons. These findings will support further development of gene therapies for inner retinal disease or conditions where production of soluble gene products is required.

## Results

As previous studies had mainly reported transfection of Müller glia in the inner retina, we were particularly interested in investigating the tropism of the LNPs beyond Müller cells. We commenced this investigation in cultured dissociated retinal cells. By this approach, LNPs would have direct access to all cell types of the inner retina and thus, LNP tropism beyond the influence of any anatomical barriers could be studied. We therefore transfected the cultured dissociated retinal cells with cmRNA*^mCherry^*m^1^Ψ_1.0-_m^5^C_1.0_–LNPs and 24h later, mCherry expression was evaluated. Indeed, also in dissociated cells, the largest portion of transfected cells was immunopositive for GFAP and had a drawn-out structure typical for Müller cells **(Figure 1A)**. Among the GFAP^+^ cells, we also encountered some that were rather star-shaped, thus likely representing astrocytes **(Figure 1B)**. Most interestingly, among the transfected cells we also repeatedly found GFAP immunonegative cells with a neuron-like morphology **(Figure 1C, asterisk**), whereof one specifically had a typical bipolar-cell morphology and was additionally immunopositive for the ON-Biopolar cell marker Protein Kinase C alpha (PKCα) **(Figure 1C, arrowhead)**. The rest of these neuron-shaped cells were immunonegative for PKCα (and all other markers tested). Some mCherry^+^ cells also exhibited morphological similarities with macrophages (**Figure 1D, arrowhead**). Thus, at least in the absence of anatomical barriers, LNPs are able to transfect diverse retinal cell types, including neurons.

**Figure 1:**
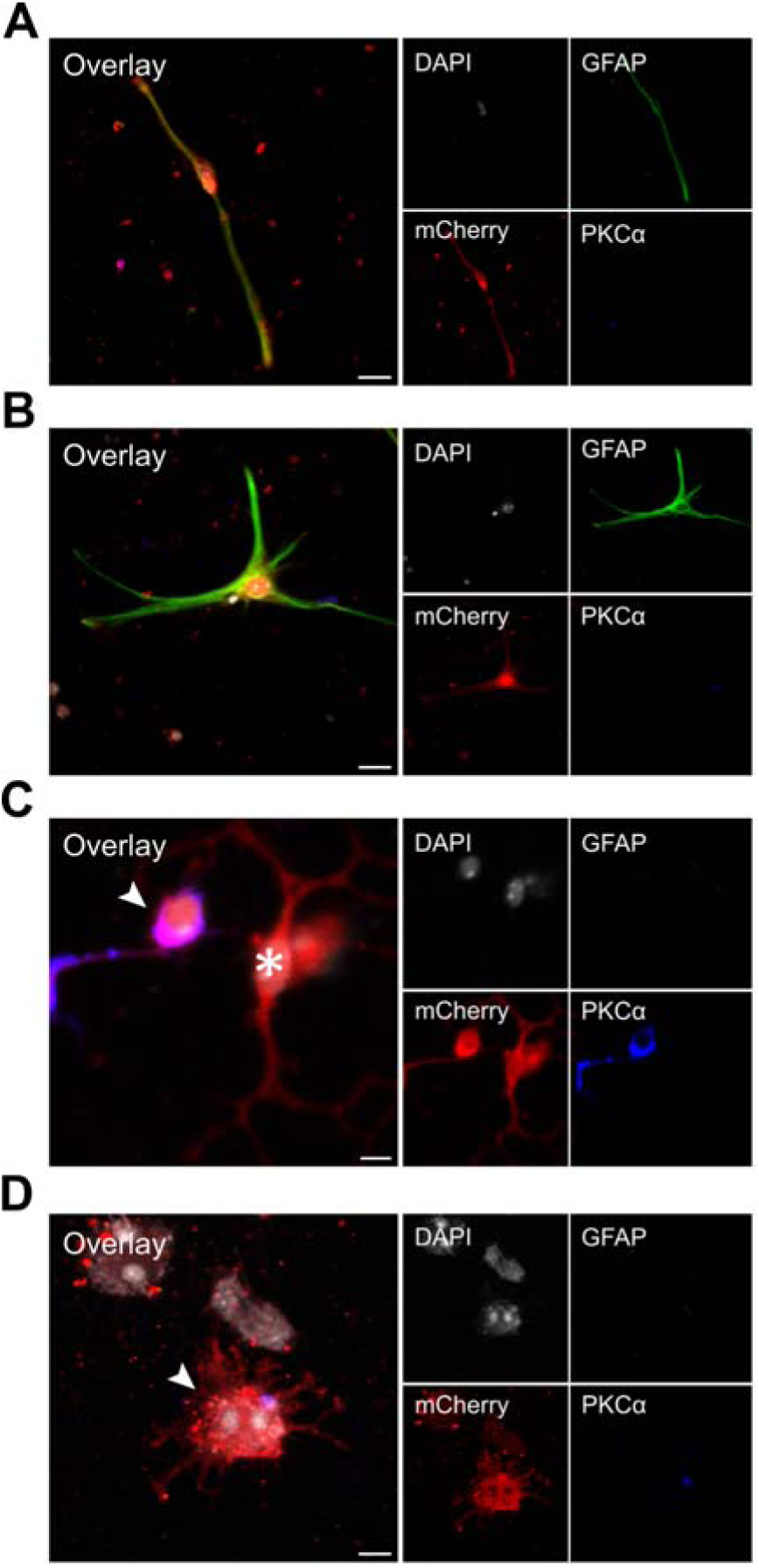
Transfection of cultured dissociated retinal cells using cmRNA*^mcherry^*m^1^Ψ_1.0_m^5^C_1.0_-LNP. Representative confocal micrographs of dissociated primary retinal cells from rd1 retina-degenerate mice, fixed and immunostained 24 hours post-transfection for mCherry (red), the Müller glia and astrocyte marker GFAP (green) and the rod bipolar cell marker PKCα (blue). Counterstain: DAPI (gray). (A) Müller glia. (B) Astrocyte. (C) Rod bipolar cell and other neuron. (D) Macrophage. Scale bar: A, B, D= 10 µm, C= 5 µm.

We next explored which cell types could be transfected by LNPs in the intact retina. As a starting point, we applied cmRNA*^mCherry^*m^1^Ψ_1.0-_m^5^C_1.0_–LNPs to retinal explants from retina-degenerate rd1 mice that were cultured ganglion cell-side down. Thus, the application of the LNPs to the surface of the explant would resemble a subretinal route of delivery. Transfection was performed immediately after taking the explants into culture (hereafter referred to as acute transfection). Twenty-four hours later the retinas were fixed and assessed for mCherry immunofluorescence. Robust mCherry expression could be observed throughout all treated explants (**Figure 2A**). On average, we observed 34.89 [±4.01] mCherry^+^ cells per field of view (FoV) (**Figure 2C**). Notably, there was a substantial variability in transfection yield within individual retinas (**Supplementary Figure 1**), which is consistent with previous observations^12^. To find out how many of the transfected cells would be Müller glia, we assessed co-incidence of the mCherry signal with Glial Fibrillary Acidic Protein (GFAP) immunoreactivity within an individual cell (**Figure 2B, upper panel, arrowhead** indicating the overlap of mCherry^+^ cell with Müller glia process; **asterisk** pointing to the corresponding soma). Indeed, 26.33 [±3.76] cells/FOV were GFAP^+^/mCherry^+^ (**Figure 2C**) while the others showed no evidence of GFAP signal neither in their soma nor their extensions (**Figure 2B, lower panel, asterisk**).

**Figure 2:**
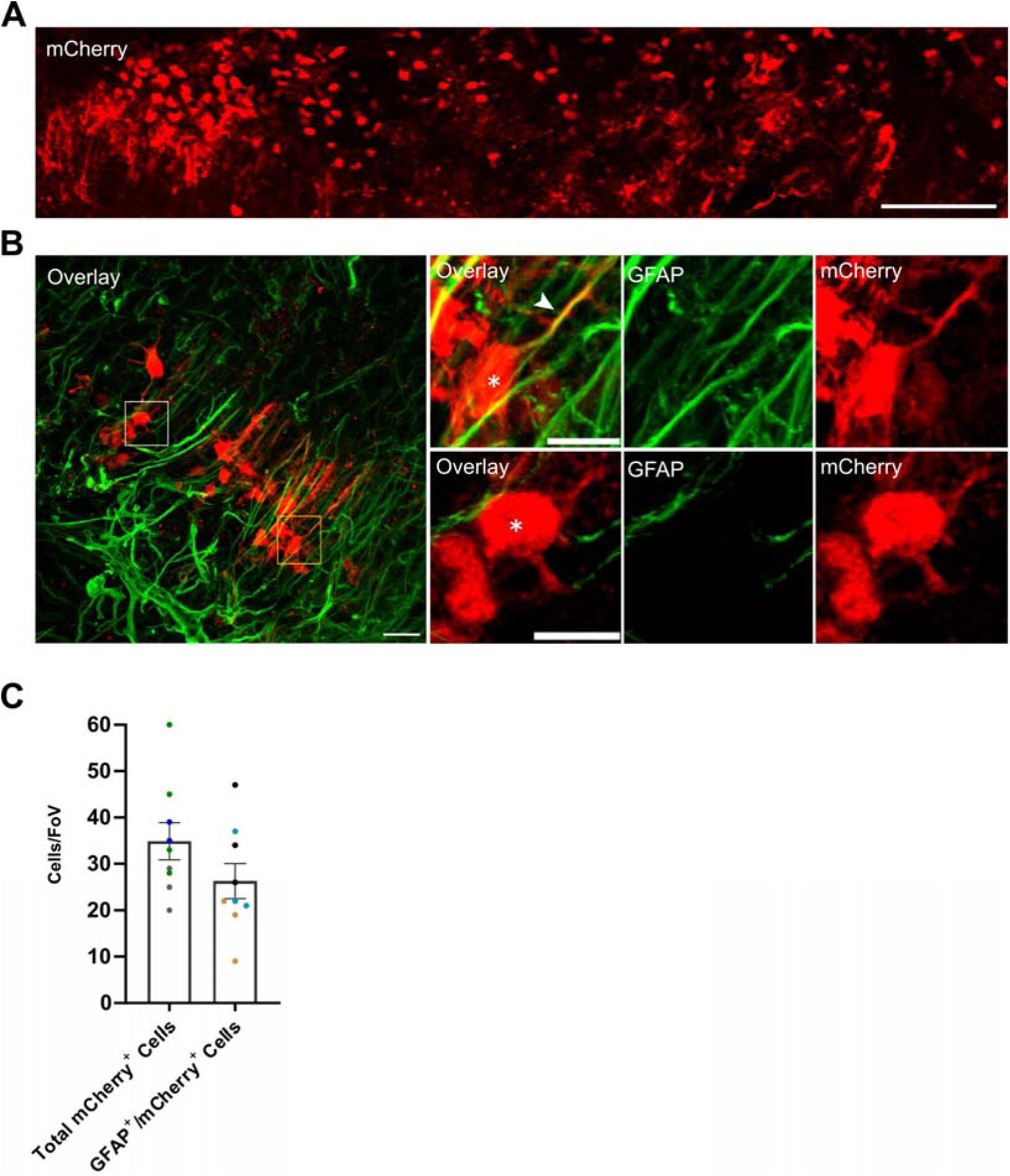
Transfection pattern in retinal explants from retina-degenerate eyes using cmRNA*^mCherry^*m^1^Ψ_1.0_m^5^C_1.0_-LNP. Retinal explants from rd1 retina-degenerate mice transfected by quasi-subretinal delivery immediately after explantation (150□ng/µL). (A) Stitched composite image was generated from multiple overlapping micrographs to visualize the full extent of transfection across the explant. (B) Exemplary micrographs of explants immunostained for mCherry (red) and GFAP (green). (C) Number of total mCherry^+^ and GFAP^+^/mCherry^+^ cells per field of view (FoV) assessed 24h post-transfection. Note that GFAP does not stain the somata of Müller glia but rather their processes, hence, cells were classified as GFAP^+^ based on their processes. Dots represent data acquired from an individual FoV, colours represent individual biological replicates (3 per condition). A FoV represents a single confocal micrograph (212.55 µm x 212.55 µm). Scale bar: A= 150 µm, B= 20 µm, B upper and lower panel= 10 µm.

To confirm that our observations were not biased by potential acute cell stress artefacts induced by the explantation process, we repeated these assessments in retinas kept in culture for two days before transfection (denoted as late transfection, **Supplementary Figure 2A-C**). Except for a slightly, albeit statistically significant, lower proportion of GFAP^-^ cells (late: 5.88 [±0.64], acute: 8.56 [±0.88] GFAP^-^/mCherry^+^ cells/FOV; *p*<0.05, **Supplementary Figure 2D**) transfection yield was largely similar. Therefore, in all subsequent experiments we transfected the retinas immediately after explantation.

We next moved on to analyse the impact of the route of LNP delivery (intravitreal vs subretinal) as well as potential differences in transfection yield between degenerate and non-degenerate healthy retinas. By far the largest number of mCherry positive cells was observed after quasi-subretinal administration in degenerate explants. With 32.44 [±4.16] cells/FOV, this was significantly higher than after intravitreal administrations (13.44 [±1.79] cells/FOV, *p*<0.0001; **Figure 3A**). A similar trend was observed in healthy explants, though transfection yields were overall lower. Subretinal delivery again outperformed intravitreal delivery (intravitreal: 4.58 [±0.80], subretinal: 7.90 [±0.82] cells/FOV; *p*<0.05, **Figure 3B**). Given the reduced transfection in healthy retina and the known structural differences in the inner limiting membrane (ILM) between healthy and degenerate tissue^18,19^, we hypothesized that an intact ILM might hinder intravitreal transfection upon intravitreal delivery. Therefore, we performed an additional experiment where we mechanically removed the inner limiting membrane (ILM) from non-degenerate healthy explants (**Supplementary Figure 3**). Indeed, after ILM peeling, quasi-intravitreal LNP administration yielded 8.52 [±0.92] mCherry^+^cells/FOV, which is significantly more than what was achieved without peeling (*p*<0.01; **Figure 3B**).

**Figure 3:**
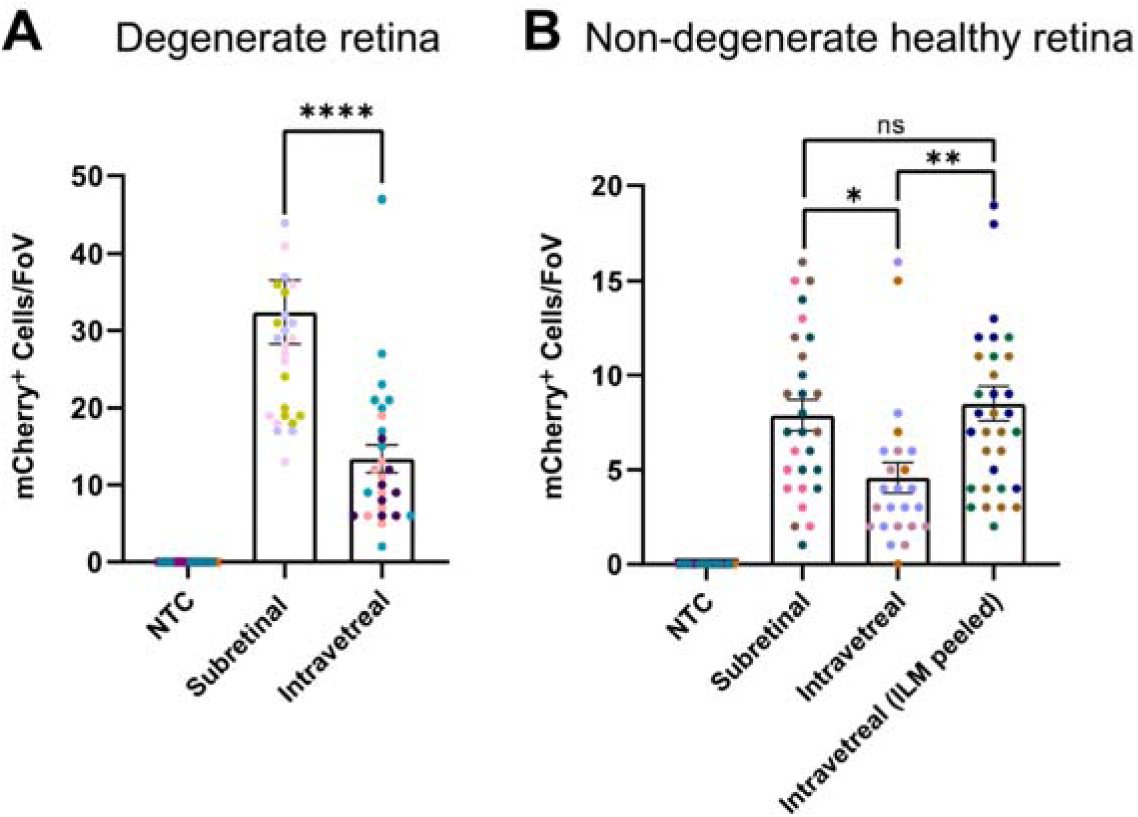
Transfection yield in degenerate and non-degenerate healthy retinal explants. Quantification of mCherry^+^ cells in retinal explants from degenerate (rd1, C3H; A) and non-degenerate healthy (C57BL/6J; B) mice following quasi-subretinal or intravitreal delivery of LNPs (150□ng/µL). Non-treated control (NTC) retinas were used as controls. Dots represent data acquired from an individual FoV, colours represent individual biological replicates (3 per condition). A FoV represents a single confocal micrograph (212.55 µm x 212.55 µm). To enhance visualization in panel 3A, the y-axis was truncated. Two outlier values from subretinal administration fall beyond the axis limit and are not displayed, but were still included in the statistical analysis. Data in panel 3A were analyzed using the Mann-Whitney test, and data in panel 3B using one-way ANOVA followed by Tukey’s HSD post hoc test. Unless otherwise indicated, data are presented as mean ± SEM. Statistical significance is indicated as ns (non-significant), * *p* < 0.05, ** *p* < 0.01, *** *p* < 0.001 or **** *p* < 0.0001.

To characterize the non-glial targets in the intact retina more in detail, we performed an in-depth morphological analysis of the GFAP^-^ mCherry^+^ cells in the cultured explants. Following quasi-intravitreal delivery, we observed mCherry expression in structures resembling blood vessels, thus in cells representing pericytes or endothelial cells **(Figure 4).** Similar observations could not be made following quasi-subretinal delivery. Moreover, consistent with our observations in dissociated retinal cells, also in the intact retina we could observe mCherry^+^ cells with neuron-like morphology. We further assessed these by neurite tracing on confocal volume scans. We found that the somata of these cells were typically located to the outer portion of the inner nuclear layer (INL) and were consistently immunonegative for PKCα. By localization and morphology, in particular, the horizontal orientation of their neurites, we postulate that these are Horizontal cells (**Figure 5A and B, and Supplementary Figure 4A and B**). Additionally in degenerate retinas, we also detected a small number of mCherry^+^ cells distributed within the ganglion cell layer (GCL) of the retina (**Figure 5C**).

**Figure 4:**
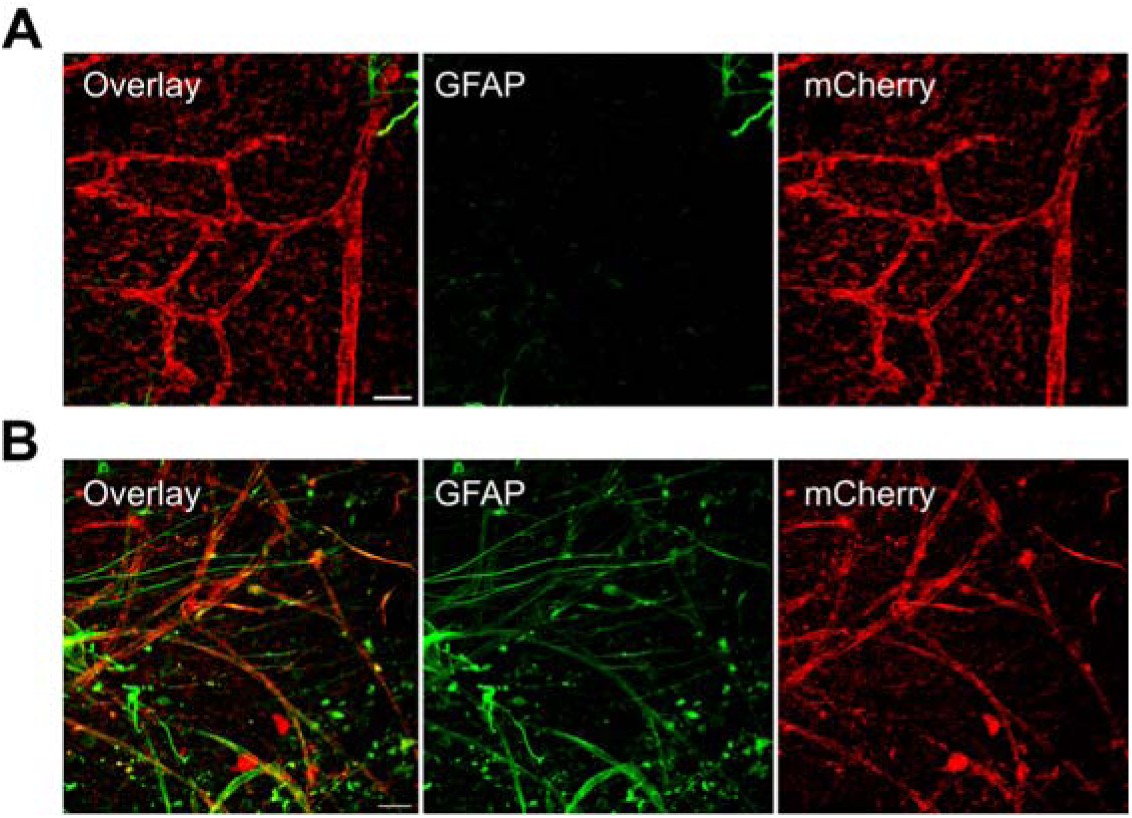
cmRNA*^mCherry^*m^1^Ψ_1.0_m^5^C_1.0_-LNP transfects blood vessels. Micrographs acquired from non-degenerate healthy retinal explants from C57BL/6J mice that were subjected to quasi-intravitreal delivery of LNPs (150□ng/µL). Immunostaining for mCherry (red) showed the transfection of vessel-like structures in (A) non-degenerate healthy retina (ILM-peeled) and (B) in non-degenerate healthy retina after quasi-intravitreal administration. Note that occasionally one can observe GFAP-immunoreactivity (green) closely aligning (but not completely overlapping) with these structures (presumably representing astrocytes sheathing those vessels). Scale bar: 20 µm. For better visualization background noise was removed using despeckling and local averaging (median) filters. Data obtained from 3 independent biological replicates. Similar observations were made on degenerate rd1 retinas following quasi-intravitreal delivery (not shown).

**Figure 5:**
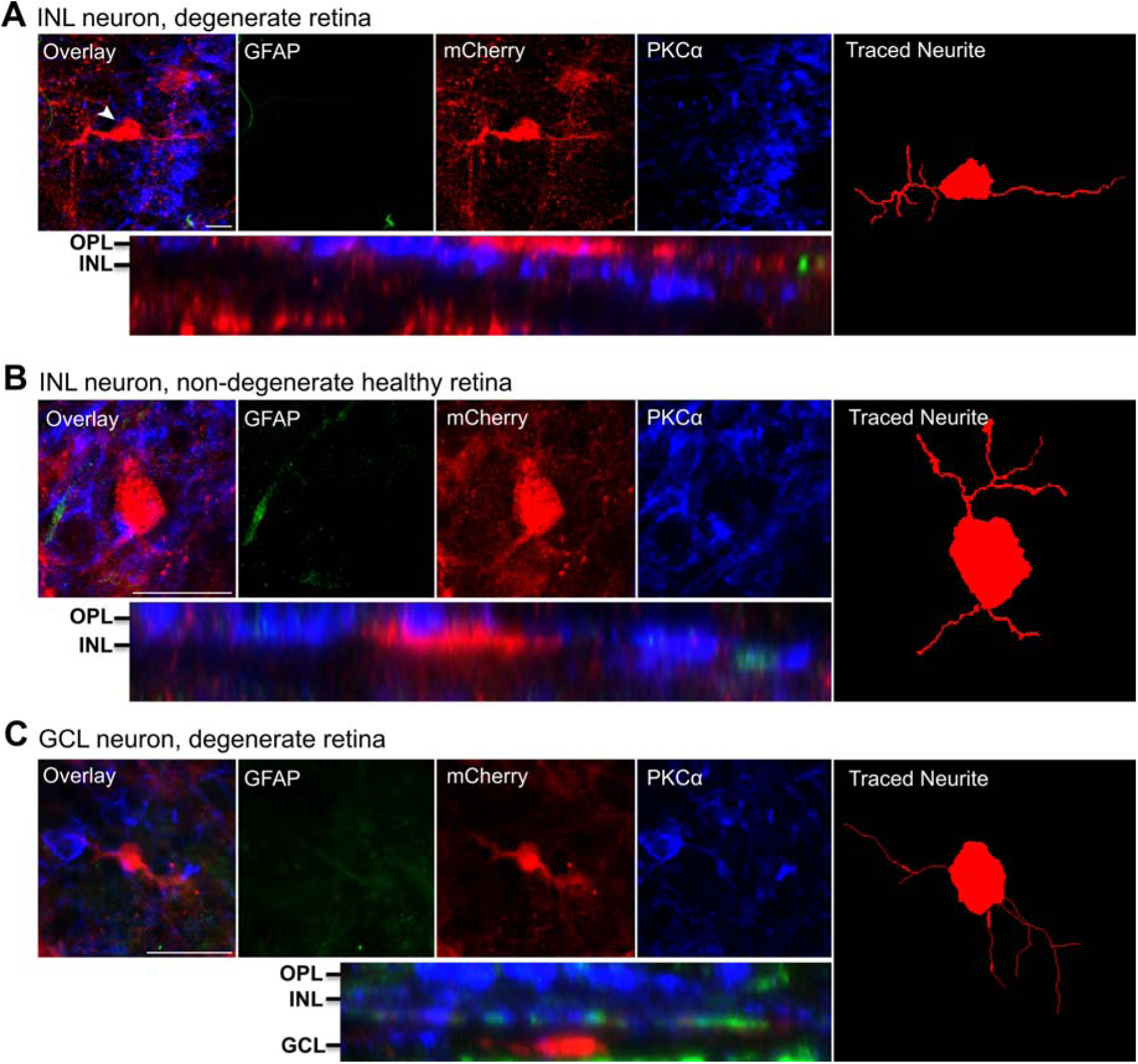
cmRNA*^mCherry^*m^1^Ψ_1.0_m^5^C_1.0_-LNP transfects retinal neurons. Exemplary high-resolution volume scans from degenerate (rd1, C3h; A and C) retinal explants 24 hours post intravitreal and subretinal transfection, respectively and non-degenerate healthy (C57BL/6J; B) retinal explants 24 hours following subretinal transfection. Retinal explants were triple-labelled with mCherry (red), GFAP (green) and PKCα (blue). Corresponding orthogonal representations of the cells are shown below. Arrowhead represents the associated traced neurite in the right panel. OPL: outer plexiform layer, INL: inner nuclear layer, GCL: ganglion cell layer. Scale bar: A, B= 20 µm.

While LNPs have been generally reported to exert little toxicity, LNPs themselves or the genetic material they deliver may trigger inflammatory or other stress responses in the retina^17,20^. A sensitive marker for retinal stress is Müller glia activation, which is associated with an increase in GFAP expression in these cells^21^. We therefore assessed differences in the levels of GFAP immunoreactivity. In degenerate explants, GFAP levels were markedly and significantly increased following intravitreal administration (*p*<0.0001) as well as subretinal administration (*p*<0.01) compared to non-transfected control (NTC) explants **(Figure 6A and B, and Supplementary Table 1)**. In non-degenerate healthy explants, intravitreal administration without prior ILM peeling only led to a small and non-significant (*p*<0.507) increase in GFAP signal as compared to NTC controls. Upon subretinal administration, by comparison, this increase was robust and statistically significant (*p*<0.001), yet still substantially lower than what had been observed in the degenerate retinas. In ILM-peeled healthy explants following intravitreal administration, GFAP signal was substantially higher than in retinas without prior peeling (*p*<0.01) **(Figure 6C and D, and Supplementary Table 1)**.

**Figure 6:**
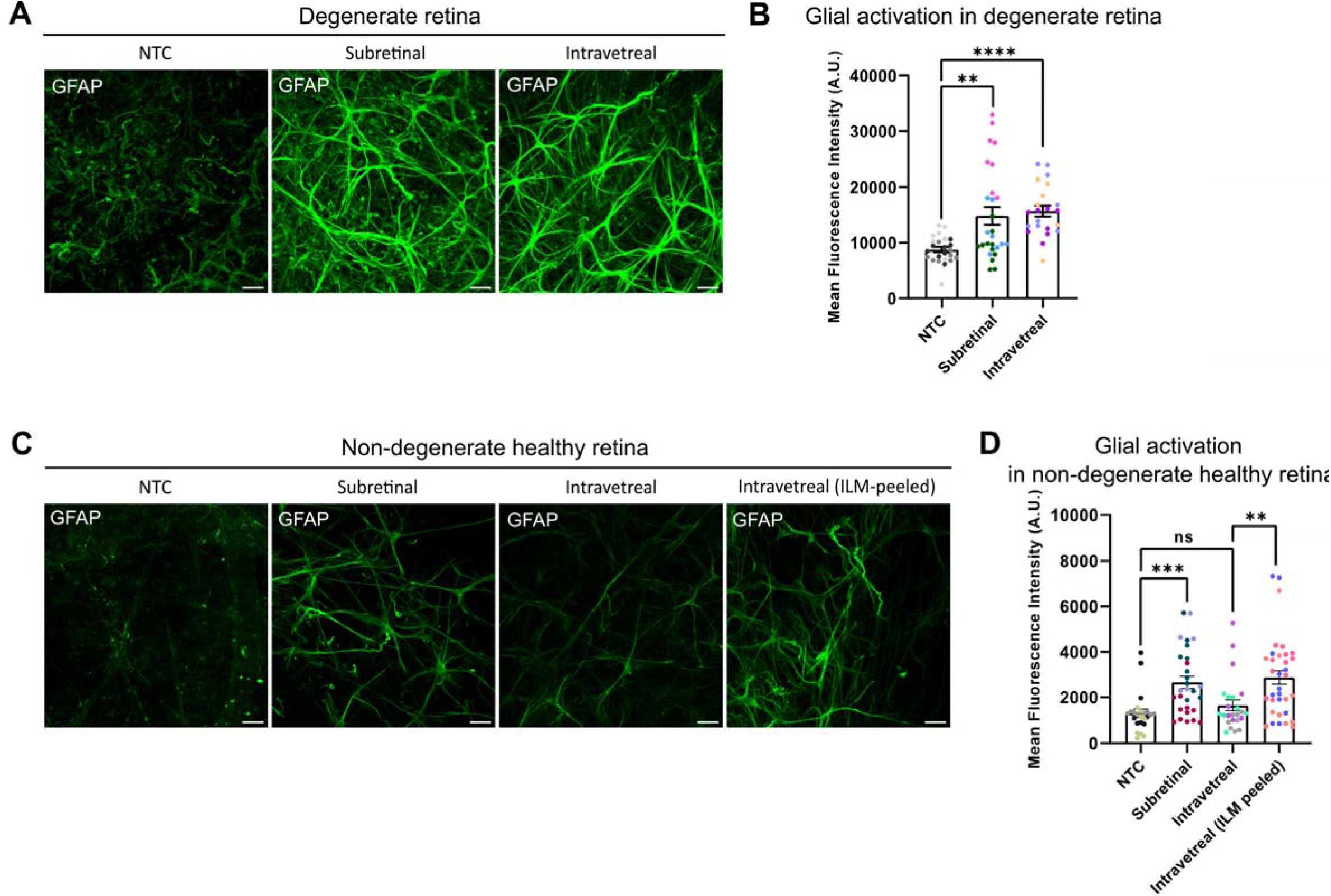
cmRNA*^mCherry^*m^1^Ψ_1.0_m^5^C_1.0_-LNP mediated glial activation across retinal explants. Representative confocal micrographs of GFAP immunoreactivity from (A) degenerate (rd1, C3H) and (C) non-degenerate healthy (C57BL/6J) explants after 24 hours post transfection following quasi-intravitreal and quasi-subretinal administration. Quantification of GFAP signal intensity (mean fluorescence intensity, MFI in arbitrary units [A.U.]) in (B) degenerate and (D) non-degenerate healthy retina. Respective non-treated control (NTC) groups were used as controls. Scale bar: 20 µm. Dots represent data acquired from an individual FoV, colours represent individual biological replicates (3 per condition). A FoV represents a single confocal micrograph (212.55 µm x 212.55 µm). Statistical significance in panels 6B and 6D was assessed using one-way ANOVA followed by Dunnett’s test. In panel 6D, comparisons between intravitreal administration in retinas and intravitreal administration in ILM-peeled retinas were analyzed using the Mann-Whitney test. Data are expressed as mean ± SEM, unless stated otherwise. *p*-values are reported as ns (non-significant), * *p* < 0.05, ** *p* < 0.01, *** *p* < 0.001 or **** *p* < 0.0001.

## Discussion

In the present study, we systematically characterized the cellular tropism and transfection efficiency of lipid nanoparticles delivering chemically modified mRNA in murine retinal explants and dissociated retinal cell cultures. We found that conventional LNPs transfect a broader range of retinal cell types than previously recognized including neurons and vascular cells. Quantitative analysis in explants revealed that quasi-subretinal delivery was significantly more efficient than quasi-intravitreal administration, although removal of the inner limiting membrane markedly enhanced intravitreal delivery efficacy. Furthermore, we observed that overall transfection efficiency was higher in degenerate rd1 retinas than in non-degenerate controls.

To analyse the effect of the route of delivery on LNP transfection efficacy we utilized murine retinal explants as a model system. Culturing them with the ganglion cell side attached to the cell culture insert enabled a quasi*-*subretinal route for LNP delivery while the reverse orientation (i.e., outer retina towards culture insert) allowed for quasi-intravitreal LNP delivery. We found a substantially higher transfection yield with subretinal delivery as compared to intravitreal delivery. Few studies have previously assessed the ability of LNPs to transfect inner retinal neurons using either an intravitreal^12^ or subretinal route of delivery^13,16^. Yet, direct comparisons between the two approaches are limited^11,15^ and widely descriptive. While most authors reported at least some degree of inner retinal cell (specifically Müller glia) transfection regardless of the delivery route chosen, Ryals et al. explicitly state that they did not observe any inner retinal transfection upon subretinal LNP delivery^11^. In this regard, the quantitative data we provide here are helpful in setting an objective basis for future analysis and it is particularly relevant to note that in our experience, subretinal delivery has been substantially more efficient than intravitreal delivery, despite similar local LNP concentrations, unless the ILM was removed.

The reasons for the higher transfection yield we observed upon subretinal as compared to intravitreal LNP delivery are overall unclear. We hypothesize that the ILM imposes a structural barrier due to both, pore size and its negative charge, obstructing particle transmission^22^. This concept is supported by our observation that transfection efficiency of intravitreally applied LNPs improved after mechanical ILM removal. Clearly, from the subretinal side, the outer limiting membrane (OLM) imposes a somewhat similar diffusion barrier. Yet, as opposed to the actual basement membrane nature of the ILM, the OLM is made up of the tight junctions between photoreceptors and Müller cell feet^23^. Thus, while the OLM imposes a diffusion barrier, it would not necessarily hinder particle uptake into Müller cells. In fact, there are several reported examples of material uptake into Müller cells from their apical site have been reported^24,25^. It should be noted that in our cultured explants, RPE cells were mostly absent. In an intact, non-degenerate healthy eye, these cells heavily phagocytose and may therefore scavenge LNPs that could otherwise transfect cells of the neuroretina. This impact of RPE cells might well explain why no inner retinal transfection was observed with subretinal LNP delivery in previous studies^11^.

In practice, subretinal delivery may be preferred when mRNA therapy would be a one-off treatment, as this allows for easy achievement of high local LNP concentrations, while the intravitreal route is less invasive and also possible in outpatient settings. In this regard, alternative methods for ILM removal, like enzymatic digestion^26^ or photochemical destruction^27^ may prove helpful to increase transfection yields.

We also observed substantially higher transfection efficiency in explants from retina degenerate (rd1) mice; An observation that was particularly pronounced upon subretinal delivery (**Figure 3**). This finding is of relevant given that most applications of LNP-mediated mRNA delivery to the retina are in diseased retinas. Clearly, the course of retinal degeneration is rapid in the rd1 mouse, and it is unclear to what extent our observations can be generalised to other retinal diseases. However, changes to the limiting membranes, in particular the OLM, are observed relatively early in the disease course and resemble those seen in other types of retinal degeneration^28^. In this regard, the rd1 model that we have studied herein represents an informative starting point for further studies on more specific disease models.

Previous studies have reported Müller glia as the only cells of the inner retina that can be transfected by LNPs^11,13,16,17^. Only Chambers et al. additionally observed the transfection of macrophages^12^. Herein, we have explored the capacity of LNPs to transfect inner retinal cells beyond these classically phagocytosing cells. On cultured dissociated retinal cells, where the anatomical barriers of the intact retina are absent, we observed that indeed Müller glia, and macrophages are indeed targets of LNPs, but also astrocytes and neurons can be transfected. Subsequent experiments on cultured intact retinas overall confirmed this observation in a more native setting and additionally showed that also endothelial cells and/or pericytes were among those transfected. However, we did not systematically assess the relative transfection rates for the individual cell types. In the intact retina, however, we found that the transfected cells are GFAP^+^ (i.e. Müller Glia or astrocytes), by visual assessment, the vast majority of these being Müller cells. Interestingly, amongst the remaining GFAP immunonegative cells, both neurons and vascular cells were observed and thus reveal to be feasible targets for LNPs. Hence, tailoring LNP formulations to the intended target cell type could facilitate the development of mRNA therapeutics with precise cell-type specificity. The necessary modifications for both, improved endothelial and neuronal targeting have been studied extensively outside the eye^29–31^ and could be adopted for ocular delivery.

Although by characterizing distinctive morphological features and using immunohistochemical markers we were able to coarsely infer cell identity, a more fine-granular classification of the transfected cell types, particularly among neurons, would be desirable. Future studies employing single-cell RNA sequencing, a broader set of immunohistochemical markers on retinal cryosections or multicolour flow cytometry will be helpful to comprehensively map the tropism of LNPs within the retinal cellular landscape.

Reactive Müller cell gliosis is a universal response mechanism of the retina to stress and activated Müller glia characteristically exhibit increased GFAP immunoreactivity^21^. We have therefore used GFAP signal intensity as a non-specific but sensitive marker with which to measure retinal stress in response to LNP administration. Indeed, we observed Müller cell activation following LNP administration for most conditions studied. Only after intravitreal LNP delivery to non-degenerate healthy retinas, no significant increase in GFAP signal could be observed. Since ILM-peeled explants exhibited a statistically significant increase in GFAP signal compared to non-peeled explants following intravitreal delivery of LNPs, it is possible that the ILM protects the retina from LNP-induced stress while also representing a relative obstacle for transfection. It is unclear, whether this effect is simply related to the lower intraretinal effective dose of LNPs or if there is any sort of additional protective effect (e.g., by charge filtering). In this context, previous studies have focused on assessing immune activation in response to LNP delivery and find no evidence for immune activation upon intravitreal delivery^12,17,20^ and some upon subretinal delivery^16,17^. Of note, Müller glia activation may also occur independent of immune activation and thus represents a complementary way of assessing the retina response to LNP-mediated transfection. Nevertheless, more differentiated immune profiling should be performed using the established RNA ImmunoGenic assay^32^ should be performed and minimizing the immunogenicity of the delivered RNA cargo could be achieved by further chemical nucleoside modifications, codon optimization, and/or incorporation of miR-142-3p binding sites into the 3′ UTR to restrict expression in immunogenic cell types^7,33,34^.

From a translational perspective, the key observation made in this study is that inner retinal cell types beyond the phagocytosing cells – including neurons – can be transfected using LNPs. In particular, the transfection of neurons is of relevance as it indicates that treatment of hereditary retinal disorders that have their primary cause in the inner retina, like several forms of Congenital Stationary Night Blindness (CSNB), by LNP-mediated mRNA delivery is generally feasible. Bipolar cells, the main therapeutic target in many CSNB subtypes, have proven difficult to transduce with adeno-associated virus (AAV) vectors^35^. LNPs would therefore provide an attractive alternative. Moreover, the 4.7 kb packaging ceiling of AAVs (promoter included) precludes delivery of large genes like TRPM1 – a major CSNB locus – or of the multi-component gene-editing systems required for in situ correction^6,36^. LNPs, by contrast, have a much higher payload and can encapsulate all gene-editing components in a single particle. With regard to gene-editing, the non-perpetual nature of LNP-mediated transfection is an additional advantage to minimize the risk of off-target effects^6^. Certainly, this will require further tweaking of LNP tropism in favour of neuronal transfection – wherefor recent advances made in transfecting brain neurons could serve as template^31^. In this regard, it is worth noting that in our studies on dissociated retinal cells transfected (PKCα^+^) ON-bipolar cells were particularly rare, suggesting that there might be cell-type specific challenges that need to be addressed.

Beyond these requirements for cell-type specific transfection, our results demonstrate that efficient transfection of Müller cells can be achieved using conventional LNPs with components that are all already employed in clinical settings^37^ and an intravitreal route of delivery (enabling repeated administration e.g., in an outpatient setting), efficient transfection of at least of Müller cells can be achieved. Thus, this approach could prospectively be exploited in regenerative applications, where the aim is to reprogram Müller glia into neurons ^38,39^ or where soluble gene products are needed. Such situations include the production of VEGF-receptor fragments to treat exudative retinal disease^40^, complement inhibitors to halt atrophic AMD^1^ or neurotrophic factors for Glaucoma or Macular Telangiectasia Type II^41,42^. In this context, pharmacokinetics could be adjusted using depot systems and materials^43^ or self-amplifying mRNA formats^44^.

In addition to these translational considerations, our finding that LNPs can be efficiently used for gene delivery to cultured retinal explants is of practical relevance in laboratory research settings. Compared to alternative virus-mediated gene delivery, LNP-mediated expression occurs much faster, accelerating experiments and reducing the time it takes for cultured organs to drift.

### Conclusions

Our findings demonstrate that conventional lipid nanoparticles (LNPs) can efficiently transfect a range of retinal cell types beyond Müller glia, including neurons and vascular cells, particularly in degenerate retinal tissue. Subretinal delivery consistently outperforms intravitreal administration in terms of transfection efficiency, although removal of the inner limiting membrane can partially overcome this limitation. These results highlight the potential of LNPs as a flexible and scalable platform for gene therapies targeting inner retinal diseases, especially where transient expression or large payloads are required.

## Materials and Methods

### mRNA and LNP formulation

mRNA encoding the fluorescent protein mCherry encapsulated in lipid nanoparticles were purchased from VectorBuilder (Chicago, USA). To generate cmRNA*^mCherry^*m^1^Ψ_1.0_m^5^C_1.0_, uridine-tri-phosphate (UTP) and cytidine-tri-phosphate (CTP) were completely substituted by N^1^-methylpseudouridine (m^1^Ψ) and 5-methylcytosine (m^5^C), respectively. LNP was formulated with (6Z,9Z,28Z,31Z)-Heptatriaconta-6,9,28,31-tetraen-19-yl 4-(dimethylamino)butanoate (Dlin-MC3-DMA), 1,2-Distearoyl-sn-glycero-3-phosphocholine (DSPC), Cholesterol, and 1,2-dimyristoyl-rac-glycero-3-methoxypolyethylene glycol-2000 (DMG-PEG_2000_) in a ratio of 50:10:38.5:1.5. Subsequent quality assurance assays were performed by the manufacturer.

Two batches of LNP were produced and characterized by dynamic light scattering. Particle sizes were 73.7 nm and 101.1 nm, with polydispersity indices of 0.072 and 0.070, respectively. Zeta potentials were -4.07 mV and -9.82 mV, respectively.

### Animals

C57BL/6J and C3H/HeOuJ mice were purchased from Charles River (Sulzfeld, Germany). All the mice were housed in a specific pathogen-free animal facility on a 12 h light-dark cycle with food and water available *ad libitum*. Tissue collection was carried out in accordance with federal law and institutional guidelines. For tissue collection, mice were decapitated following deep isoflurane (Baxter, Deerfield, Illinois, United States) anesthesia and retinas were extracted and immediately transferred to CO_2_-independent medium (Thermo Fisher Scientific, Waltham, MA, United States) as previously described^45^.

### Primary retinal cell culture and transfection

Retinas were dissociated using the Papain Dissociation System (Worthington Biochemical Corporation, Lakewood, NJ, USA) with 250µL of papain, 150µL of ovomucoid and 15µL of DNase solution per retina. Centrifugation was performed at 124 x g for 5 minutes to separate cells from debris. Retinal cells were then transferred into culture medium (Neurobasal-A, Thermo Fisher Scientific supplemented with: 10% Dulbecco’s Modified Eagle Medium, Thermo Fisher Scientific; 2.5% B-27 plus supplement, Thermo Fisher Scientific; 5µg/mL forskolin, Sigma-Aldrich, St. Louis, USA; 10ng/mL human ciliary neurotrophic factor, Miltenyi Biotec, Bergisch Gladbach, Germany) and seeded on Poly-D-Lysine and Laminin coated coverslip (Corning, New York, USA). Cells were transfected with cmRNA*^mCherry^*m^1^Ψ_1.0-_m^5^C_1.0_–LNP at a final concentration of 0.7 ng/µL. 24h after transfection, cells were fixed with 4% paraformaldehyde (PFA, Thermo Fisher Scientific) and stored in PBS at 4°C until further use.

### Retinal explant preparation and transfection

Retinal explants were prepared following the method by Devoldere et al. with minor modifications^15^. After extraction, retinas were transferred onto semi-permeable PTFE membrane cell culture inserts (Merck, New Jersey, USA) either with photoreceptor side or the retinal ganglion cell side facing upwards and fixed to the insert by applying negative pressure. These two orientations of the explant with either the ganglion or photoreceptor side facing upwards would allow mimicking an intravitreal or subretinal route of delivery, respectively^46^. Culturing medium (Neurobasal-A supplemented with 1% B-27, 1% Penicillin-streptomycin, 0.5% L-Glutamine, 2.5% fetal bovine serum) was added below the inserts, after which the explants were cultured at 37°C. The explants were transfected either immediately after being taken into culture or after culturing for two days using cmRNA*^mCherry^*m^1^Ψ_1.0-_m^5^C_1.0_–LNP at a final concentration of 150 ng/µL. 24h post-transfection, retinal explants were fixed with 4% PFA and stored at 4°C until immunostaining.

### Removal of the inner limiting membrane

Before transferal onto the cell culture inserts, the internal limiting membrane (ILM) was stained using 0.4% trypan blue (Thermo Fisher Scientific) for 1-2 minutes. Excess trypan blue was removed by gently shaking the retina inside the preparation dish. Fine forceps (Fine Science Tools, Heidelberg, Germany) were used to remove the ILM under a dissecting microscope.

### Immunocytochemistry and Immunohistochemistry

Dissociated retinal cells were permeabilized with 0.1% Triton-X in PBS (0.1% PBSTX) and blocked with 5% goat serum. Primary antibody incubation was performed in 2.5% goat serum in PBS for 2 hours and secondary antibody incubation for 1h. All steps were carried out at room temperature (RT). Stained cells were stored in PBS at 4°C.

The immunostaining of cultured retinal explants was performed as described earlier^47^. The following primary antibodies were used: Anti-GFAP (1:1000 dilution, catalog no. ab7260, Abcam, RRID: AB_305808), anti-PKCα (1:500, sc-8393, Santa Cruz, RRID: AB_628142), anti-mCherry (1: 500, ab205402, Abcam, RRID: AB_2722769) and anti-Laminin (LAMA1) (1:800, L9392, Sigma-Aldrich, RRID: AB_477163). The secondary antibodies used were: goat anti-rabbit 488 IgG (H+L; A11008, Thermo Fisher Scientific) goat anti-mouse 647 IgG1 (γ1; A21240, Thermo Fisher Scientific) and goat anti-chicken 568 IgY (H+L; ab175477, Abcam), all used at a concentration of 1:1000.

### Image acquisition and analysis

Confocal micrographs were acquired using a LSM 710 confocal microscope (Zeiss, Oberkochen, Germany) as previously described^47^. For explants, volume scans covering the entire depth of the inner retina were acquired using a 40x oil immersion objective. The laser settings were kept constant for all images recorded per experimental series. In experiments aimed at quantifying the number of mCherry-positive cells or fluorescence intensity in cultured explants, adjacent volume scans were acquired across the entire explant – from one edge to the other – to capture a representative portion of the explant in its full depth. Care was taken to minimize any overlap between adjacent scans. For imaging of dissociated cells, a 20x water objective was used.

Image analysis was performed using ImageJ/Fiji (National Institute of Health, Bethesda, MD)^48^. When stated, Neurite tracing of mCherry^+^ cells was performed semi-automatically using the Simple Neurite Tracer (SNT) plugin^49^. Overlapping tiles were stitched in three dimensions with the pairwise stitching plugin^50^ and the stitched image was z-projected with maximum intensity. Post-processing was limited to global brightness and contrast adjustments, downscaling, cropping and selection of relevant fluorescent channels when appropriate.

### Statistical analysis

Statistical analysis was performed using GraphPad Prism version 8 (GraphPad Software, San Diego, CA, USA). Data are presented as mean ± SEM. For comparison between two groups, either a Mann-Whitney U test or an unpaired t test was performed as appropriate. Unless stated otherwise, comparisons involving more than two groups were analyzed using one-way ANOVA; when the ANOVA was significant, pairwise differences were tested with Tukey’s HSD. Where comparing to a single control condition, Dunnett’s test was used instead of Tukey’s HSD. Statistical significance was denoted as \**p* < 0.05, \*\**p* < 0.01, \*\*\**p* < 0.001, \*\*\*\**p* < 0.0001.

## Declarations

### Author Contributions

Participated in research design: SB, ML, VR

Conducted experiments: SB, PS, EC, NX

Performed data analysis: SB

Wrote or contributed to the writing of the manuscript: SB, EC, NX, VR, ML

### Data Availability

The datasets generated during and/or analysed during the current study are available from the corresponding author on reasonable request.

### Funding

Supported by Grants from the University Medical Centre Giessen und Marburg to ML (UKGM 15/2020) and Deutsche Förderprogramm für Augenheilkunde of Bayer Vital GmbH to SB and ML.

### Declaration of Interest

This work has been supported by Bayer Vital GmbH. Bayer had no role in the design of the experiments.

### Ethics approval

This study did not involve human participants. Animal work was performed with approval of the relevant authorities and in accordance with the institutional Ethics Guidelines of Animal Care. Further details are provided in the Methods section.

### Consent to publish

Not applicable.

## Supplementary information

**Supplementary table 1:**
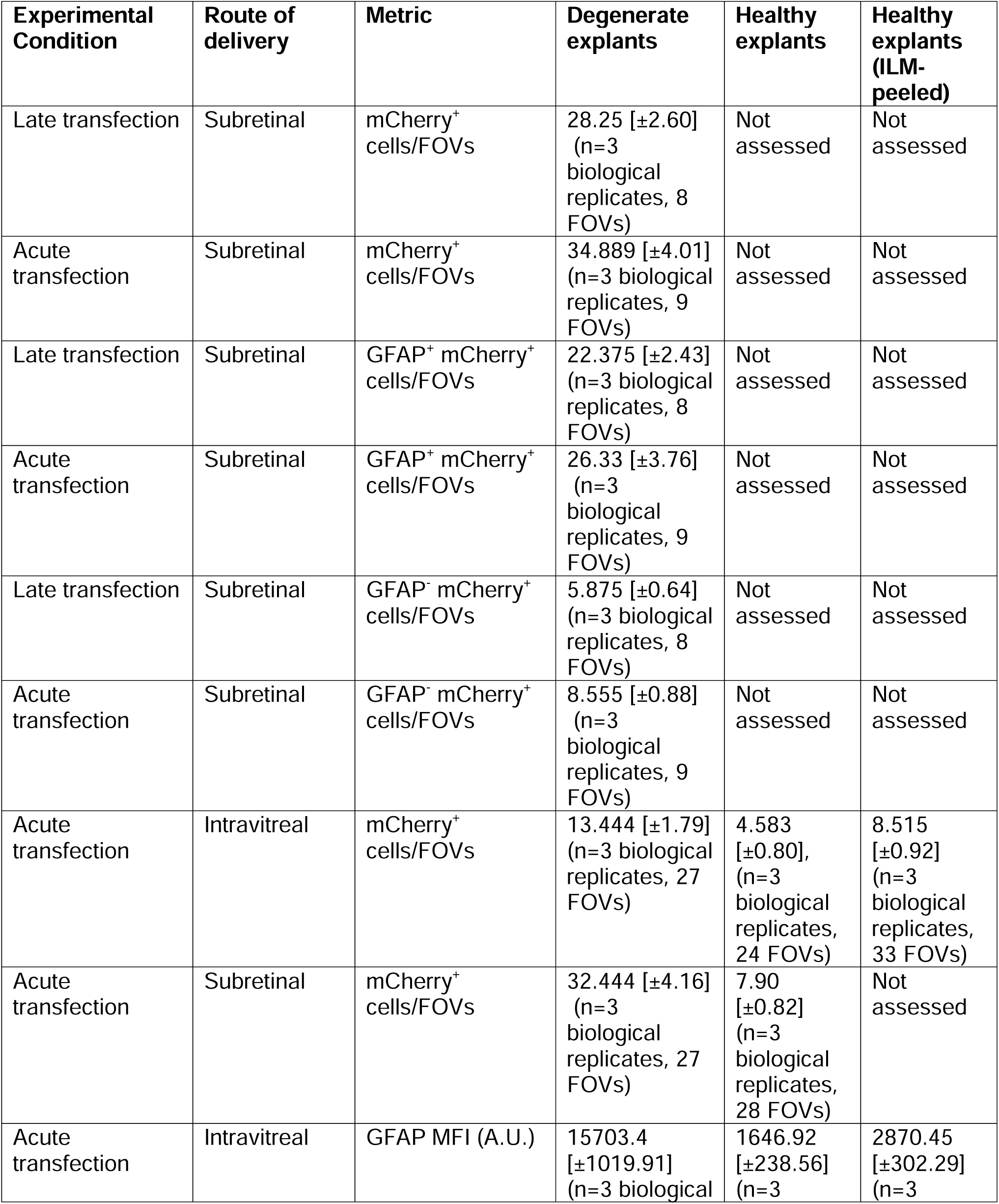
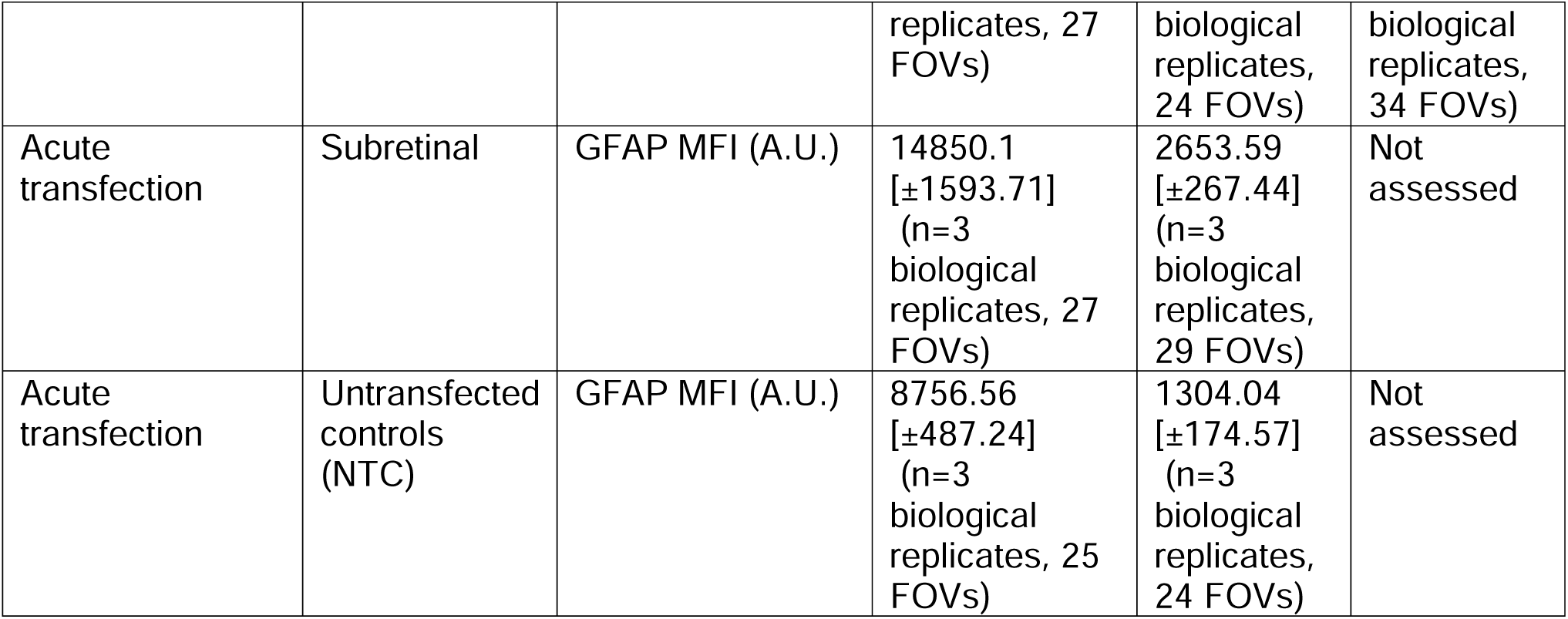
Summary of reported metrics in this study.

### Supplementary Figures

**Supplementary figure 1:**
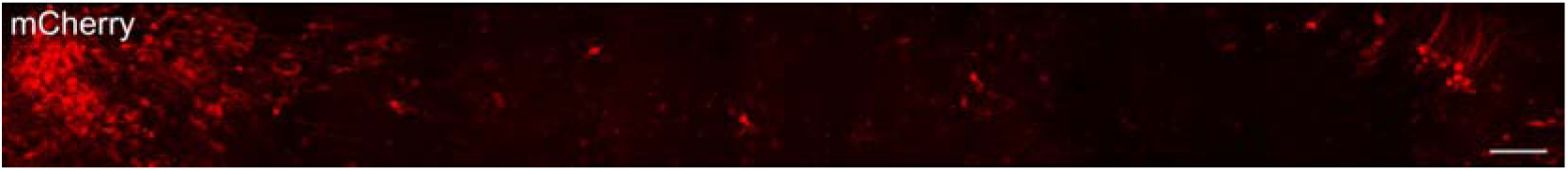
Comprehensive visualization of cmRNA transfected region in retina. Retinal explants from rd1 retina-degenerate mice transfected by quasi-subretinal delivery (150□ng/µL). Multiple overlapping micrographs were stitched into a composite image to allow the assessment of transfection throughout the retinal explant. Scale bar: 150 µm.

**Supplementary figure 2:**
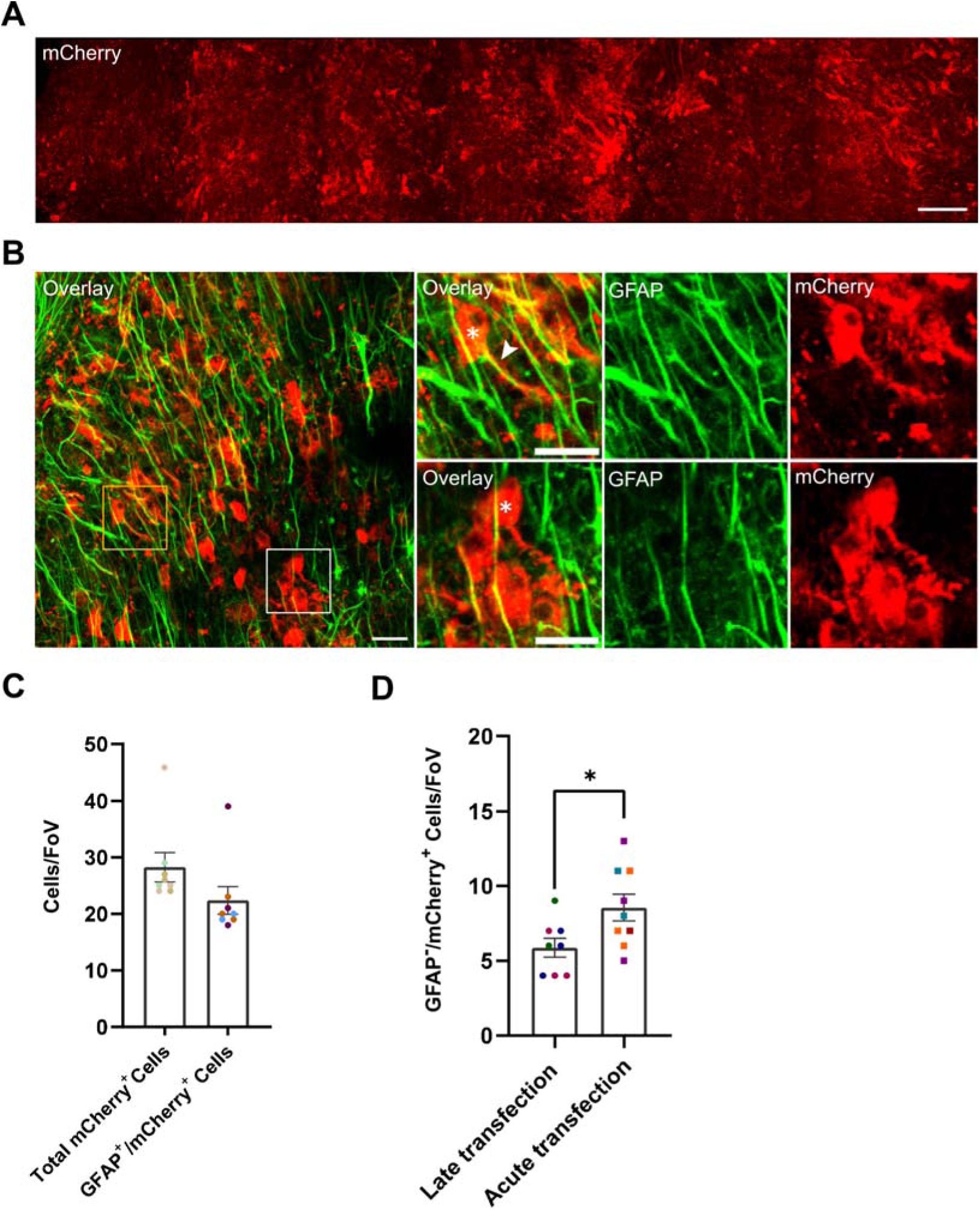
Late transfection in degenerate retinal explants using cmRNA*^mcherry^*m^1^Ψ_1.0_m^5^C_1.0_-LNP. Degenerate retinal explants from rd1 mice transfected by quasi-subretinal delivery using LNPa two days after explantation. (A) Stitched composite image was generated from multiple overlapping micrographs to visualize the full extent of transfection across the explant. (B) Representative micrographs of explants immunostained for mCherry (red) and GFAP (green). Arrowhead represents the overlapping transfected cells and Muller glia process. Asterisk depicts a GFAP^-^/mCherry^+^ cell (C) Number of total mCherry^+^ cells and subset of GFAP^+^/mCherry^+^ cells per field of view (FoV) after late transfection, assessed 24 hours post-transfection. (D) Quantification of GFAP^-^/mCherry^+^ cells per FoV, representing the complementary subset of the total mCherry^+^ cells after late and acute transfection. Note that GFAP does not stain the somata of Müller glia but rather their processes, hence, cells were graded as GFAP^+^ based on their processes. Scale bar: A= 150 µm, B= 20 µm, B upper and lower panel= 10 µm. Data obtained from 3 independent biological replicates per condition. Statistical significance of the data in panel 2D was performed using an unpaired t test. Unless specified otherwise, data are shown as mean ± SEM. *p*-values are reported as ns (non-significant), * *p* < 0.05, ** *p* < 0.01, *** *p* < 0.001 or **** *p* < 0.0001.

**Supplementary figure 3:**
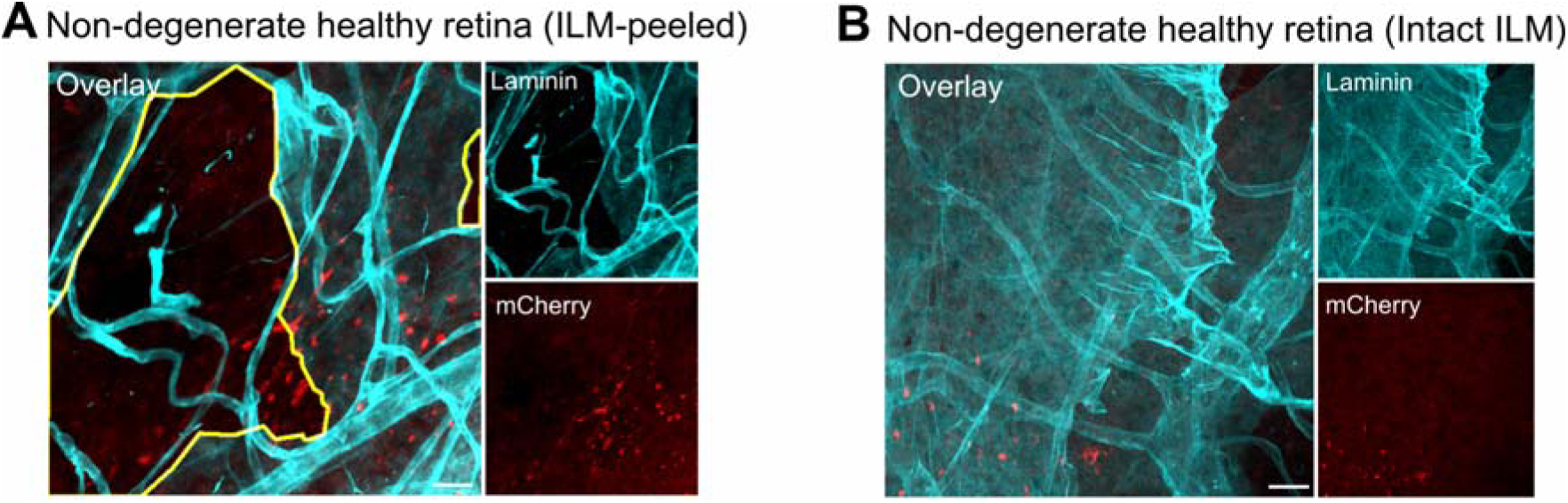
ILM-peeling promotes cellular uptake of cmRNA*^mcherry^*m^1^Ψ_1.0_m^5^C_1.0_-LNP. ILM was partially removed during explantation of non-degenerate healthy explants (C57BL/6J). A representative confocal micrograph of (A) partially removed ILM and (B) intact ILM. Flat-mount retinal explants were double-stained with ILM marker laminin (cyan) and mCherry (red). ILM-peeled areas are highlighted (yellow). Scale bar: 20 µm.

**Supplementary figure 4:**
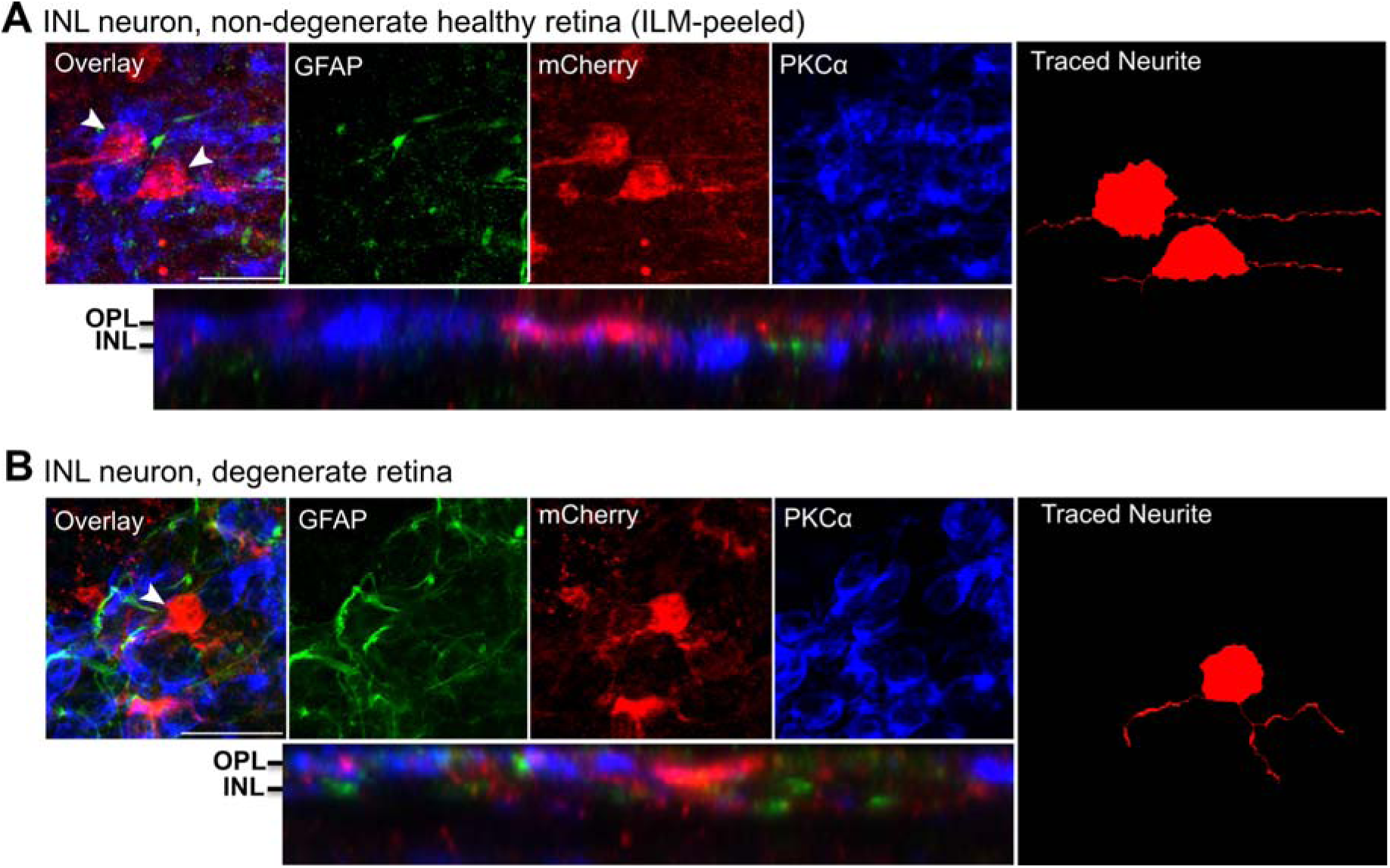
cmRNA*^mcherry^*m^1^Ψ_1.0_m^5^C_1.0_-LNP transfects non-degenerate healthy and degenerate retinas. Exemplary image from a high-resolution volume scan of (A) non-degenerate healthy explants (C57BL/6J) following intravitreal (ILM-peeled) and (B) degenerate explants (rd1, C3H) after subretinal administration. Flat-mount staining was performed using GFAP (green), mCherry (red) and PKCα (blue). Left panel: z-projection of the neuron-like structure with maximum intensity, with corresponding montage. Right panel: corresponding traced neurites. Arrowhead represents the associated traced neurite in the right panel. OPL: outer plexiform layer, INL: inner nuclear layer. Scale bar: 20 µm.

